# MutaStruct: Revisiting Benchmarks for Single-Site Mutation-Induced Structural Change Prediction

**DOI:** 10.64898/2026.10.21.760837

**Authors:** Tan-Phat Huynh, Dang-Manh Ho, Viet Thanh Duy Nguyen, Quang-Hien Kha, Ngoc-Thac Pham, Kuei-Hung Lai, Truong-Son Hy

## Abstract

Predicting the structural consequences of single-site mutations is fundamental to understanding protein function, disease mechanisms, and protein engineering. However, existing benchmarks for mutation-induced structural change prediction are often affected by duplicate mutation events, highly similar structural pairs, and experimental-state confounders, which can inflate reported performance. We introduce MutaStruct, a curated benchmark of experimentally observed single-mutation protein structure pairs designed for rigorous evaluation of mutation-induced structural changes. MutaStruct combines residue-level sequence validation, coordinate-level quality control, context-aware filtering, event-level deduplication, and leakage-aware data splitting to improve dataset reliability and evaluation rigor. We benchmark sequence-only structure prediction, physics-based refinement, and learning-based refinement methods together with a Copy-WT diagnostic control. Copy-WT remains highly competitive on backbone-based metrics, while method rankings change substantially across local and mutation-site measures. These results show that overall structural fidelity does not necessarily imply accurate modeling of mutation-associated structural change, underscoring the need for further methodological development beyond current paradigms. The benchmark dataset is currently undergoing further curation and expansion and will be released in a future journal-version update. The codebase, trained models, and evaluation pipeline are publicly available at https://github.com/HySonLab/MutaStruct.

## 1 Introduction

Single-site amino acid substitutions are a fundamental driver of protein evolution, and understanding how they reshape protein structure is central to connecting genotype to phenotype. A single residue change can alter a protein’s fold, stability, or interaction surface, with consequences ranging from loss of function to disease. Many single-site mutations, for example, are directly involved in cancer and other disorders [4, 6, 7, 9]. Accurately characterizing the structural consequences of such mutations is therefore a prerequisite for both basic mechanistic biology and applied efforts such as variant interpretation and protein engineering.

Experimental structure determination remains the gold standard for resolving these structural changes. However, techniques such as X-ray crystallography are low-throughput and costly, and can only be applied to a small fraction of the mutations of interest. Computational structure prediction has partially filled this gap: methods such as AlphaFold [10] now predict the overall tertiary structure of most proteins with remarkable accuracy. However, several studies have shown that AlphaFold and related structure predictors are not sensitive to the subtle structural perturbations caused by a single amino acid substitution. Given a wild-type sequence and its single-mutation variant, these models frequently predict near-identical structures regardless of the true structural outcome [3, 15]. This insensitivity limits the usefulness of general-purpose structure predictors for mutation-specific structural analysis, and motivates the development of methods specialized for this task.

Recent work has begun to address this gap directly. PreMut [13] was introduced as the first deep learning approach to predict the mutant tertiary structure directly from a wild-type structure, reporting substantial improvements over AlphaFold and establishing this template-conditioned refinement setting as a promising direction beyond prior mutation-effect methods, which largely predict scalar outcomes such as stability change or pathogenicity rather than structure itself [18, 20]. However, we argue that the benchmarks used to evaluate this task have limitations that obscure how much genuine mutation sensitivity has actually been achieved. Because most atoms in a single-site mutant remain structurally unchanged, a trivial baseline that simply copies the wild-type structure is likely to score competitively on standard global metrics such as TM-score and GDT-HA, suggesting that these benchmarks may reward preservation of the unchanged structure more than accurate modeling of the mutation’s effect. Existing train/test splits also rely on sequence-identity clustering, which does not guarantee structural novelty, and wild-type-mutant pairs curated from sequence matching alone can conflate the mutation’s true effect with unrelated experimental differences such as ligand occupancy or crystallographic conditions. Together, these issues mean that the performance gains reported may not reliably reflect improved sensitivity to mutation-induced structural change.

To address these limitations, we introduce MutaStruct, a revised benchmark for single-site mutation structure prediction. It (1) curates mutation-controlled, context-validated structure pairs to reduce confounding from non-mutation experimental differences, (2) applies structure-aware, leakage-controlled dataset splitting, and (3) stratifies evaluation by the magnitude of true structural change and reports local, mutation-centered metrics alongside global structure scores. Using this benchmark, we re-evaluate three broad categories of methods: Structure Prediction Models, which predict a mutant structure from sequence alone; Physics-Based Refinement Models, which perturb a wild-type template via energy-function minimization; and Learning-Based Refinement Models, which learn the structural correction directly from data given a wild-type template and mutation, alongside a copy-baseline control. We find that Copy-WT remains highly competitive in backbone-based metrics, while method rankings change substantially when overall local distance agreement and mutation-site side-chain accuracy are considered. These results show that high absolute structural similarity does not by itself demonstrate accurate modeling of mutation-associated structural change, and motivate evaluation that explicitly separates backbone preservation from local mutation-site accuracy.

## 2 Benchmark Construction

We construct a benchmark for learning and evaluating experimentally observed structural responses to single amino-acid substitutions. A central consideration is that a mutation event and a pair of deposited structures are not equivalent: two structures differing by one residue may also differ in ligand occupancy, metal coordination, crystallographic environment, or experimental state, and the same mutation is often deposited across independent crystal forms. As illustrated in Fig. 1, our curation pipeline therefore combines sequence-level mutation discovery, reference–variant pair construction, context matching, coordinate-level quality control, and leakage-aware partitioning to produce a benchmark in which each retained pair reflects a well-defined, context-controlled mutation event.

**Figure 1:**
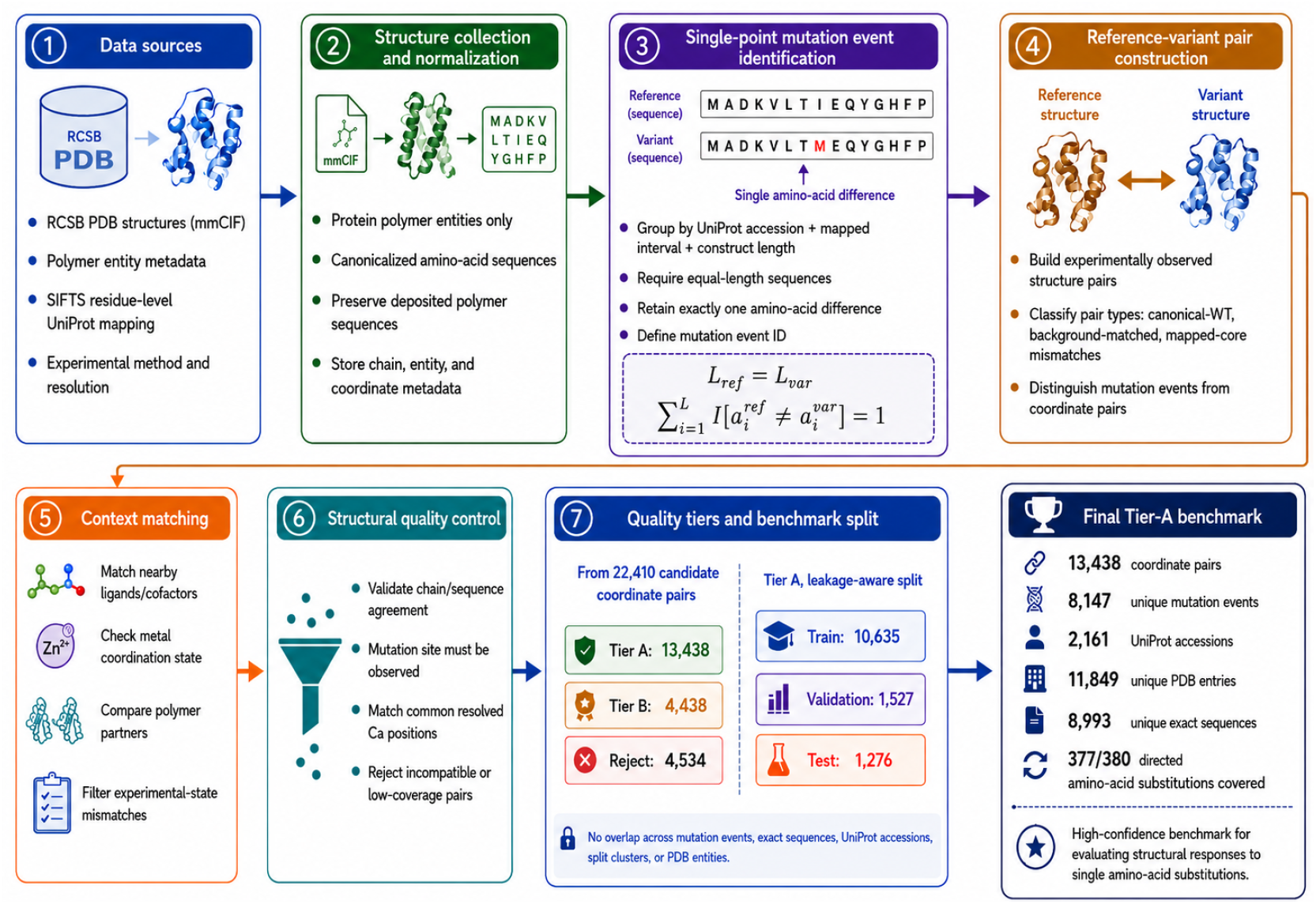
Overview of the MutaStruct dataset construction pipeline. Experimentally determined protein polymer entities are collected from the PDB and linked to UniProt through residue-level SIFTS mappings. Single-substitution mutation events are identified from equal-length sequence pairs, followed by construction of experimentally observed reference–variant structure pairs. Candidate pairs are subsequently filtered for experimental-context compatibility and structural quality. High-confidence Tier-A pairs are finally partitioned using a leakage-aware splitting procedure that prevents overlap across mutation events, exact sequences, UniProt accessions, sequence clusters, and PDB polymer entities.

Briefly, protein polymer entities are collected from the PDB and linked to UniProt via residue-level SIFTS mappings. We identify single-substitution mutation events by comparing equal-length sequence pairs, and assign each one a canonical event identifier so that repeated depositions of the same mutation are not treated as independent events. For each mutation event, candidate reference–variant structure pairs are then filtered in two ways: first, for experimental-context compatibility (matching ligand and metal state, experimental method, and resolution), to rule out structural differences caused by lab conditions rather than the mutation itself; second, for coordinate-level quality at the mutation site, to ensure the region of interest is actually resolved and usable. The resulting high-confidence (Tier A) pairs are partitioned into training, validation, and test sets using a leakage-aware procedure, which prevents any overlap across partitions at the level of mutation events, exact sequences, UniProt accessions, sequence clusters, and PDB polymer entities. The complete details of each step are provided in the Appendix A.

The resulting Tier-A benchmark contains 13,438 coordinate pairs that span 8,147 mutation events, 2,161 UniProt accessions, and 377 of 380 possible directed substitutions, with no cross-partition overlap at any of the above levels. The coordinate quality is high (median common C*α* coverage 98.8%), and while structural changes are predominantly subtle (median global C*α* RMSD 0.270 Å), the benchmark retains a long tail of larger and locally concentrated rearrangements.

### 3 Experimental Results

We evaluated MutaStruct on the Tier-A test split using five complementary metrics that span global, local, and mutation-response structural agreement: Global C*α* RMSD, all-atom lDDT, Local C*α* RMSD_8Å_, mutation-site side-chain RMSD, and MR-RMSD_8Å_. For the RMSD-based metrics, we first remove global rigid-body differences using a Kabsch superposition over mutually represented C*α* atoms; this global transform is reused for Local C*α* RMSD_8Å_ and mutation-site side-chain RMSD without additional local realignment, while lDDT requires no superposition since it is defined from internal interatomic distances. MR-RMSD_8Å_ instead aligns the predicted and experimental variant structures independently to the reference structure before comparing their local reference-to-variant C*α* displacement fields, directly testing whether the predicted structural response improves upon a no-change hypothesis rather than whether the predicted structure is merely close to the mutant in absolute terms. We report this combination because global metrics alone are dominated by the majority of the structure left unchanged by a single-site mutation and can therefore reward trivial preservation of the reference structure, whereas the remaining metrics isolate agreement specifically at and around the mutation site. The full metric definitions are provided in the Appendix C.

We compare three baseline categories, chosen to span the range of paradigms currently used for mutation-induced structural modeling. *Structure Prediction* methods predict the mutant structure directly from sequence without access to the wild-type template; we include this category because it represents the general-purpose structure prediction setting most commonly used in prior work to assess mutation sensitivity, and serves as a reference point for how much a wild-type template actually contributes to accurate mutation modeling. *Physics-Based Refinement* methods instead take the wild-type structure and mutation as input and perturb it via energy-function minimization; we include this category because it represents the classical, pre-deep-learning approach to mutation-induced structural modeling, and provides a non-learned baseline against which the benefit of data-driven refinement can be assessed. *Learning-Based Refinement* methods likewise condition on the wild-type structure and mutation, but learn the structural correction directly from data rather than an explicit energy function; we include this category because it represents the emerging paradigm this benchmark is designed to stress-test, and directly evaluates whether learned models generalize beyond simply reproducing the wild-type structure. Alongside these, we report *Copy-WT*, a trivial baseline that returns the wild-type structure unmodified; we include it not as a competing method but as a diagnostic control, to quantify how much of each metric and method’s apparent performance reflects genuine mutation sensitivity rather than trivial preservation of the unchanged structure. Because Copy-WT never instantiates the mutant residue identity, it cannot produce a valid atom topology at the mutation site, so lDDT and side-chain RMSD are omitted for this baseline. Appendix B introduces each baseline method and its adaptation to the mutation-conditioned structure refinement setting used in this benchmark.

## 4 Discussion

A key observation from Table 1 is that the trivial Copy-WT baseline achieves competitive performance on Global C*α* RMSD and MR-RMSD_8Å_, comparable to or better than more sophisticated methods. This behavior is expected: for single-site mutations, the backbone conformation rarely changes significantly from the wild-type structure. Methods such as FoldX, which optimize side-chain conformations while keeping the backbone fixed, achieve similarly low C*α* RMSD values precisely because the backbone is unchanged by design. This reveals a fundamental limitation of C*α*-based metrics for evaluating mutation structure prediction: they are often dominated by the largely preserved backbone and can therefore be insensitive to localized mutation-associated changes. Mutation-site side-chain RMSD therefore provides a complementary measure of local chemical correctness at the substituted residue, which may be important for packing, binding, and molecular interactions.

**Table 1:** Performance comparison of representative protein mutation structure prediction and refinement methods on MutaStruct. ↓indicates lower is better and ↑indicates higher is better. Bold denotes the best value in each column.

| Category | Method | Global $C_\alpha$ RMSD (Å) ↓ | IDDT ↑ | Local $C_\alpha$ RMSD<sub>8Å</sub> ↓ | Mutation-site SC RMSD (Å) ↓ | MR-RMSD<sub>8Å</sub> ↓ |
| --- | --- | --- | --- | --- | --- | --- |
| Trivial Baseline | Copy-WT | 0.3528 | – | 0.3375 | – | 0.3375 |
| Structure Prediction | AlphaFold 3 [1] | 0.9758 | 0.9274 | 0.6477 | **1.2277** | 0.6480 |
|  | ESMFold [12] | 0.9223 | 0.9214 | 0.6047 | 1.2726 | 0.6052 |
| Physics-Based Refinement | FoldX [8] | **0.3527** | 0.9389 | **0.3374** | 1.4682 | **0.3374** |
|  | Rosetta [5] | 0.4519 | 0.9581 | 0.4993 | 1.3149 | 0.4935 |
| Learning-Based Refinement | PreMut (EGNN-based) [13] | 0.3718 | 0.9687 | 0.3395 | 2.4896 | 0.3395 |
|  | GeoTransformer [16] | 0.3530 | **0.9705** | 0.3378 | 1.3316 | 0.3378 |

A second observation is that method rankings change substantially depending on which aspect of the mutant structure is evaluated. In particular, high overall all-atom agreement does not necessarily imply accurate modeling of the substituted residue. PreMut achieves an lDDT of 0.9687, among the highest values in Table 1, yet its mutation-site side-chain RMSD is 2.4896 Å, substantially higher than those of the other evaluated methods. Conversely, AlphaFold 3 and ESMFold achieve considerably lower mutation-site side-chain RMSDs of 1.2277 and 1.2726 Å, respectively, despite substantially larger Global C*α* RMSD and MR-RMSD_8Å_. GeoTransformer occupies an intermediate regime, combining high lDDT with comparatively low side-chain error while remaining essentially indistinguishable from the no-change baseline on the C*α*-based response metrics. These differences highlight that backbone preservation, overall local distance agreement, and mutation-site chemical geometry are related but distinct objectives. Because the substituted residue contributes only a small fraction of the atom pairs entering a global lDDT score, a prediction can preserve the overall local distance network while still placing the mutant side chain incorrectly. Accordingly, no single metric provides a complete description of mutation-structure prediction quality.

The contrasting behavior of sequence-only and reference-conditioned methods further illustrates this metric dependence. AlphaFold 3 and ESMFold achieve the lowest mutation-site side-chain RMSDs despite larger backbone errors. Their MR-RMSD_8Å_ values should not be interpreted solely as mutation insensitivity, because the score also includes ordinary de novo structure-prediction error.Conversely, FoldX and Rosetta benefit from starting from the experimental reference structure and therefore preserve the backbone well, but this does not necessarily translate into more accurate mutant side-chain placement. Together, these results reinforce that backbone preservation and local chemical accuracy are distinct aspects of the task.

More broadly, the challenge is to identify when a mutation requires deviation from an already strong reference-structure prior while avoiding unnecessary changes elsewhere. The strong Copy-WT baseline reflects the predominantly subtle reference–variant differences in MutaStruct, although a long tail of mutations shows larger structural responses. MutaStruct should therefore be viewed primarily as a benchmark for mutation-aware structural modeling rather than for stability, binding affinity, biological function, or pathogenicity. Its PDB-derived composition also means that it should not be assumed to represent the distribution of clinically relevant variants.

## 5 Conclusion

We presented MutaStruct, a curated benchmark for evaluating protein structural changes induced by single-site mutations. Compared with existing datasets, MutaStruct incorporates residue-level sequence validation, coordinate-level quality control, context-aware filtering, event-level deduplication, and leakage-aware data partitioning, providing a more rigorous and realistic evaluation setting for mutation-induced structural change prediction. Using MutaStruct, we systematically evaluated representative methods spanning structure prediction, physics-based refinement, and learning-based refinement.

Our results show that a trivial baseline returning the unmodified reference structure remains competitive with, or superior to, more sophisticated methods on backbone-based metrics, highlighting the strong no-change prior created by the predominantly subtle structural response to single-site substitutions. More importantly, method rankings differ substantially across evaluation axes. Methods that preserve the reference backbone well do not necessarily place the substituted side chain accurately, while high all-atom lDDT can coexist with substantial mutation-site side-chain error. Conversely, sequence-only structure predictors achieve comparatively accurate mutant side chains despite larger backbone deviations. These results indicate that backbone preservation, overall local geometric agreement, and mutation-site chemical accuracy are distinct aspects of the prediction problem, and that no evaluated method consistently performs well across all three. Mutation-induced structural change prediction therefore remains an open problem in which high absolute structural similarity should not be interpreted as evidence of mutation-specific accuracy.

We believe MutaStruct provides a useful resource for studying mutation-conditioned protein structure modeling and developing methods that better capture local structural adaptation caused by amino acid substitutions. Several extensions could further strengthen the benchmark. First, targeted evaluation of mutations exhibiting larger backbone and long-range conformational responses would provide a more sensitive test of mutation-dependent structural prediction. Second, paired wild-type and mutant predictions from sequence-only models would allow their predicted structural response to be evaluated without conflating it with ordinary de novo prediction error. Explicitly modeling experimental structural variability and alternative conformational states would further help distinguish reproducible mutation-associated changes from measurement- or state-dependent differences. Finally, extending MutaStruct beyond single-site substitutions to multi-site mutations would enable the study of non-additive structural effects relevant to protein engineering, variant analysis, and therapeutic design.

## A Benchmark Construction Details

### A.1 Data Sources

We obtained protein structures and polymer-entity metadata from the RCSB Protein Data Bank (PDB) [2] using the Search and Data APIs. We retained experimentally determined protein polymer entities together with their mmCIF coordinates, experimental method, resolution, release metadata, and entity and chain identifiers. Residue-level correspondence between PDB polymer entities and UniProt sequences was obtained from SIFTS [19]. We performed sequence comparisons at the polymer-entity level rather than the PDB-entry level because a single PDB entry can contain multiple proteins, repeated chains, nucleic acids, ligands, or cofactors. Both the deposited polymer sequence and a canonicalized standard-amino-acid sequence were retained; the latter was used for mutation discovery, while the former was preserved to identify non-standard polymer chemistry.

### A.2 Single-Point Mutation Event Identification

Candidate mutation events were identified using a sequence-first procedure. Polymer entities were grouped by UniProt accession, SIFTS-mapped interval, and construct length. Within each group, two canonicalized sequences were considered a candidate pair only when they had equal length and differed at exactly one position,

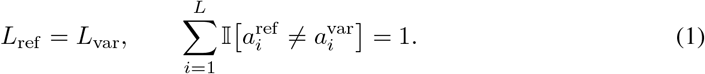

Pairs containing insertions or deletions, ambiguous residues at the substitution site, or incompatible mapped intervals were excluded from the high-confidence benchmark. Each substitution was assigned a mutation-event identifier defined by the protein background, mapped interval, mutation position, reference residue, and variant residue. This event-level representation prevents multiple experimentally deposited structures of the same mutation from being treated as independent biological mutations.

### A.3 Reference–Variant Pair Construction and Context Matching

Experimentally observed structures belonging to the same mutation event were paired only after establishing compatible sequence and structural backgrounds. We distinguish three retained pair classes. *Canonical-WT* pairs use a reference construct matching the canonical UniProt sequence over the mapped interval. *Background-matched* pairs contain a valid single substitution, but the reference should not be interpreted as a strict canonical wild type. Finally, *mapped-core mismatch* pairs (`mapped_core_length_mismatch`) contain equal-length single-substitution constructs but differ in the extent of structurally mapped or resolved coverage; these pairs are retained only if sufficient common coordinates remain for reliable comparison.

To reduce non-mutation-related structural differences, a context profile was constructed for each coordinate observation. The profile includes nearby ligand and cofactor identities, ligand-contact patterns, metal-coordination state, polymer partners, experimental method and resolution, and available experimental-state metadata. High-confidence pairs require compatible ligand presence and identities, compatible ligand-contact and metal-coordination states, compatible experimental methods, and no explicit time-resolved or photolysis mismatch. Context comparison is based primarily on the deposited asymmetric unit; biological-assembly ambiguity therefore remains a limitation of the current release.

### A.4 Quality Control and Structural Filtering

After sequence and context matching, structures were validated directly from their mmCIF coordinates. The selected polymer chains were required to agree with the expected sequences and to contain the mutation site in both structures. Backbone atoms at the mutated position were required; alternate conformations were handled deterministically, and common resolved residues were matched by polymer sequence position. Pairs with insufficient common C*α* coverage, unresolved mutation-site coordinates, incompatible experimental contexts, missing structures, or structural-processing failures were rejected.

Accepted pairs were assigned a quality tier. Tier A contains high-confidence, context-compatible pairs with strong coordinate completeness, whereas Tier B permits minor or partially tolerated limitations. Of 22,410 coordinate pairs entering the final QC stage, 13,438 were assigned Tier A, 4,438 Tier B, and 4,534 were rejected. All benchmark experiments in this work use Tier A only.

For descriptive analysis, structures were superimposed over their mutually observed C*α* positions. We calculated global C*α* RMSD, local sequence-window RMSD, spatial mutation-neighborhood RMSD, and mutation-side-chain RMSD where chemically comparable side-chain atoms were available.

### A.5 Leakage-Aware Train/Validation/Test Splitting

Random splitting at the coordinate-pair level would lead to substantial information leakage because a single mutation event may be represented by multiple experimentally deposited reference–variant structure pairs, while the same protein sequence or closely related sequences may occur in several PDB entries. We therefore performed partitioning at a grouped level rather than treating individual coordinate pairs as independent samples.

Protein sequences occurring on either side of the retained Tier-A pairs were grouped using MMseqs2 sequence clustering [17]. The splitting pipeline subsequently assigned each observation a final *split-cluster* identifier, which served as the indivisible unit of partitioning: all coordinate pairs belonging to the same split cluster were assigned to the same train, validation, or test partition. This grouping prevents alternative experimental realizations of the same mutation and sequence-linked protein backgrounds from being distributed across model-development and held-out partitions.

The resulting Tier-A benchmark contains 10,635 training pairs, 1,527 validation pairs, and 1,276 test pairs, corresponding to 6,522, 798, and 827 unique mutation events, respectively. After partitioning, we independently audited the splits for residual leakage. No mutation-event identifier, exact reference or variant sequence, UniProt accession, final split-cluster identifier, or PDB polymer entity occurred in more than one partition. The final benchmark comprises 1,621 split clusters.

### A.6 Dataset Characteristics and Structural Diversity

The final Tier-A benchmark contains 13,438 coordinate pairs representing 8,147 mutation events across 2,161 UniProt accessions, 11,849 unique PDB entries, and 8,993 exact protein sequences (Table 2). Coordinate redundancy is moderate: 69.0% of mutation events are represented by exactly one coordinate pair, the median is one pair per event, and no event contributes more than five pairs. The dataset contains 5,382 canonical-WT pairs (40.1%), 4,630 background-matched pairs (34.5%), and 3,426 mapped-core mismatch pairs (25.5%).

**Table 2:** Composition and structural characteristics of the Tier-A benchmark. RMSD and coverage are calculated over mutually observed C*α* positions.

| Characteristic | Value |
| --- | --- |
| Coordinate pairs | 13,438 |
| Unique mutation events | 8,147 |
| Unique UniProt accessions | 2,161 |
| Unique PDB entries | 11,849 |
| Unique exact sequences | 8,993 |
| Final split clusters | 1,621 |
| Canonical-WT pairs | 5,382 (40.1%) |
| Background-matched pairs | 4,630 (34.5%) |
| Mapped-core mismatch pairs | 3,426 (25.5%) |
| Median common Cα coverage | 98.8% |
| Median global Cα RMSD | 0.270 Å |
| Global Cα RMSD IQR | 0.171–0.471 Å |

Mutation coverage is broad: 377 of the 380 possible directed substitutions among the 20 standard amino acids are represented at the mutation-event level (Fig. 2a). Coordinate quality is also high. Median common C*α* sequence coverage is 98.8%, all Tier-A pairs retain at least 90% common coverage, and 82.6% retain at least 95%. X-ray diffraction accounts for 96.4% of unique structures,with a median resolution of 1.90 Å, while 3.6% were determined by electron microscopy, with a median resolution of 3.01 Å. In addition, 65.4% of Tier-A pairs contain a nearby ligand in both structures, with matched nearby ligand sets after context filtering.

**Figure 2:**
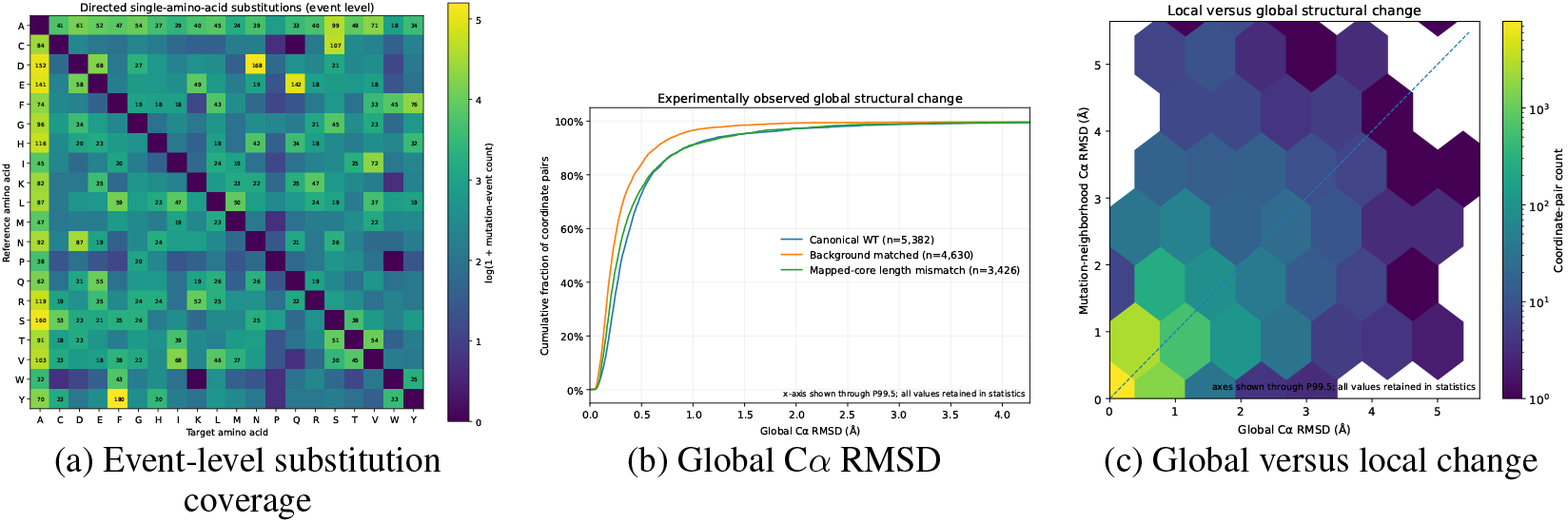
Diversity of the Tier-A mutation-structure benchmark. (a) Directed amino-acid substitutions counted once per mutation event; 377 of 380 possible directed substitutions are represented. (b)Empirical cumulative distributions of global C*α* RMSD stratified by reference–variant pair class. (c)Global C*α* RMSD versus spatial mutation-neighborhood C*α* RMSD, showing that local rearrangements can occur even when the global structure remains similar.

Experimentally observed mutation-associated structural changes are predominantly subtle but show a pronounced long tail. Pair-level global C*α* RMSD has a median of 0.270 Å (IQR 0.171–0.471 Å) and a 95th percentile of 1.221 Å. When repeated coordinate observations are collapsed by taking the median RMSD for each mutation event, the corresponding median is 0.262 Å. Overall, 93.0% of coordinate pairs have global RMSD below 1 Å, 6.1% fall between 1 and 3 Å, and 0.9% exceed 3 Å. We therefore retain RMSD as a continuous variable rather than defining these ranges as biological classes. The distributions also differ across pair classes (Fig. 2b).

Global RMSD alone does not fully describe the structural response around a substituted residue. Spatial mutation-neighborhood C*α* RMSD has a median of 0.250 Å (IQR 0.166–0.431 Å), and a subset of pairs exhibits appreciable local rearrangement despite relatively small global displacement (Fig. 2c). Thus, the benchmark spans both near-isostructural substitutions and substantially rearranged variants while maintaining high coordinate and experimental-context consistency.

### A.7 Comparison with the PreMut MutData Benchmark

The MutData benchmark introduced with PreMut [13] is the closest prior dataset for experimentally supervised single-site mutant structure prediction. MutData2022 identified mutants from PDB `SEQADV` records annotated as engineered mutations, reconstructed the corresponding wild-type sequence by reverting the annotated substitution, and searched the PDB for structures with 100% sequence identity to that sequence. This procedure yielded 36,458 possible wild-type–mutant structure pairings involving 1,094 unique mutant PDB structures. Because a mutant can be paired with several deposited structures having the same wild-type sequence, the number of reported pairs is substantially larger than the number of distinct mutant observations [13]. In contrast, MutaStruct identifies substitutions directly from polymer-entity sequence comparisons and explicitly distinguishes a biological mutation event from its potentially repeated coordinate realizations. Reference–variant pairs are additionally subjected to residue-level mapping, experimental-context matching, and coordinate-quality control before inclusion in Tier A.

Both benchmarks also account for sequence redundancy during data partitioning. PreMut clustered mutant sequences at a 30% sequence-identity threshold, producing 225 clusters in MutData2022, of which 182, 21, and 22 were assigned to training, validation, and test sets, respectively. In addition, it introduced the temporally separated MutData2023 test set, containing 59 mutants in 10 clusters with less than 30% sequence identity to MutData2022 [13]. MutaStruct complements this sequence-level control with event- and structure-entity-level constraints: complete split clusters are kept within a single partition, and the final partitions are independently audited for overlap in mutation events, exact sequences, UniProt accessions, split clusters, and PDB polymer entities. Thus, the principal difference is not simply dataset size, but the unit of representation and curation: MutData enumerates possible sequence-matched wild-type–mutant pairs, whereas MutaStruct is designed around experimentally comparable mutation events and their quality-controlled structural observations.

## B Baseline Methods

### B.1 Trivial Baseline

#### Copy-WT

Copy-WT is a diagnostic control that returns the wild-type structure unmodified as the predicted mutant structure, i.e., *Ŝ*_mut_ = *S*_wt_. No mutation information is used. Because the mutant residue is never instantiated, Copy-WT cannot produce a valid atom topology at the mutation site; therefore, we restrict its evaluation to C*α*-based metrics.

### B.2 Structure Prediction

#### AlphaFold 3 [1]

AlphaFold 3 is the latest generation of the AlphaFold family and the current state-of-the-art general-purpose structure predictor, extending beyond single-chain proteins to jointly model proteins, nucleic acids, ligands, and ions within a unified diffusion-based architecture. While previous works have extensively evaluated earlier AlphaFold versions for mutant protein structure prediction [3, 15], we specifically include AlphaFold 3 to provide an up-to-date assessment of whether the latest architectural advances translate to improved accuracy on mutation-induced structural changes, a setting that neither AlphaFold 3 nor its predecessors explicitly optimize for. For each test pair, we provide the mutant sequence as input and predict the mutant structure *de novo*, without access to the wild-type template.

#### ESMFold [12]

ESMFold is a widely used structure predictor that forgoes multiple sequence alignments entirely, instead predicting structure directly from a single sequence using representations from the pretrained ESM-2 protein language model. Its MSA-free design makes it substantially faster than alignment-based predictors and has led to its broad adoption as a fast, sequence-only baseline in structure and mutation-effect prediction studies, which motivates its inclusion here alongside AlphaFold 3. As with AlphaFold 3, we provide only the mutant sequence and predict the mutant structure *de novo*.

### B.3 Physics-Based Refinement

#### FoldX [8]

FoldX is one of the most widely used empirical force-field methods for estimating the structural and energetic effects of point mutations, and has served as a standard tool in protein stability and engineering studies for over a decade. Given a wild-type structure and a mutation specification, FoldX produces a mutant structure by locally repacking side chains and performing constrained energy minimization around the substituted residue on a fixed backbone, without requiring any learned or training-based component. We provide the wild-type structure and the mutation specification (position, wild-type residue, mutant residue) as input and take the FoldX-repaired structure as *Ŝ*_mut_.

#### Rosetta [11, 5]

Rosetta is a widely used physics- and knowledge-based macromolecular modeling suite that combines all-atom energy optimization with side-chain and backbone conformational sampling. We use PyRosetta 4 quarterly release 2026.29 as a target-blind, reference-structure-conditioned baseline. Given the experimental reference structure and mutation specification, we introduce the substitution with `MutateResidue`, locally repack residues within 8 Å of the mutation site, and apply non-Cartesian `FastRelax` using the `ref2015` score function while keeping the remainder of the structure fixed. We generate 5 decoys with 3 `FastRelax` repeats each and take the lowest-scoring model as *Ŝ*_mut_; experimental mutant coordinates are used only for evaluation.

### B.4 Learning-Based Refinement

Both Learning-Based Refinement baselines adopt a residual formulation of the mutation-conditioned structure refinement task: rather than predicting the mutant structure *X*_mut_ from scratch, the model predicts a displacementΔ*X* applied to the wild-type structure, *X*_mut_ = *X*_wt_ +Δ*X*. This formulation reflects a biophysical prior of the task: a single-site mutation rarely induces an entirely new fold, and the resulting structural change is typically small and spatially localized around the substituted residue. Framing the problem as residual prediction rather than absolute coordinate prediction is also a more favorable learning target, mirroring the use of displacement or offset prediction over absolute-coordinate regression in other structured-output domains, since a model need only learn a small, localized correction rather than reconstruct the full structure. We adopt this formulation for both Learning-Based Refinement baselines below.

#### PreMut [13]

PreMut was among the earliest deep learning approaches proposed for Learning-Based Refinement, and it is the method that first established the mutation-conditioned structure refinement setting used throughout this benchmark. PreMut represents the input structure as an all-atom graph, where each atom is a node connected to its *K* = 30 nearest neighbors, and initializes the mutated residue by copying all wild-type atom coordinates that have a counterpart in the mutant residue, while any side-chain atoms without a wild-type counterpart are randomly placed within 1.54 Å of the residue’s C*α* atom. This initial structure is passed through four equivariant graph convolutional layers (EGCL), using one-hot atom-type identity as the only invariant node feature, to predict the per-atom displacementΔ*X* that is added to the input structure via a residual connection to yield the final predicted mutant structure.

#### GeoTransformer [16]

Learning-Based Refinement remains a nascent category with very few existing methods directly applicable to the mutation-conditioned structure refinement setting; to our knowledge, PreMut is the only prior deep learning method designed explicitly for this task. To provide a second, architecturally distinct Learning-Based Refinement baseline, we adapt GeoTransformer, originally proposed for point cloud registration, to the mutation-conditioned refinement setting; to our knowledge, no prior work has applied this architecture to mutation-induced structure prediction. We select GeoTransformer for this adaptation because its geometric self-attention mechanism jointly encodes pairwise distances and triplet-wise angles between points, yielding representations that are robust to rotation and capture higher-order geometric relationships beyond the pairwise distance information used by EGNN-based architectures such as PreMut; we hypothesize this richer geometric context may better capture the propagation of structural change outward from a mutation site. We adopt the same residual formulation as PreMut, predictingΔ*X* rather than *X*_mut_ directly, to ensure direct comparability.

## C Evaluation Metrics

For one benchmark pair, let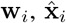, and **x**_i_ ∈ ℝ^3^ denote an atom’s coordinates in the reference, predicted mutant, and experimental mutant structures, respectively. Let *C* be the set of mutually represented and experimentally observed C*α* atoms. The global Kabsch transform is

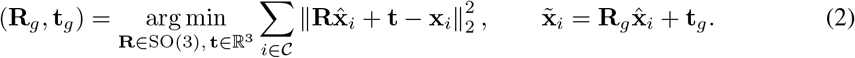

### Global C*α* RMSD

This metric measures overall backbone agreement after the global fit in Eq. (2):

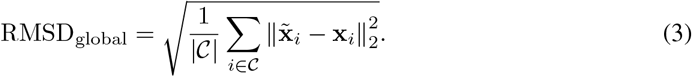

Lower values indicate better agreement.

#### All-atom lDDT

Let *P*_15_ contain unique pairs of predicted and experimentally observed heavy atoms from different residues whose distance in the experimental mutant is below 15 Å. After symmetry-equivalent atom-name correction, let 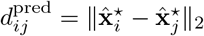 and 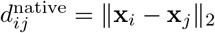, we use

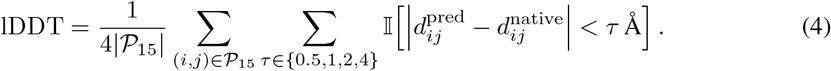

This superposition-free score lies in [0, 1], with higher values indicating better preservation of local distances [14]. Standard symmetry-equivalent side-chain atom names are treated as interchangeable before scoring.

### Local C*α* **RMSD**_8**Å**_

Let *m* be the mutation site and define the fixed reference neighborhood

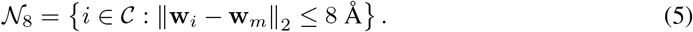

Using the same global transform as Eq. (2), rather than independently realigning the local region, we compute

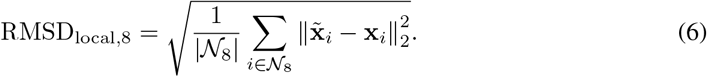

Lower values indicate more accurate mutation-neighborhood backbone geometry.

### Mutation-site side-chain RMSD

Let *S*_m_ be the set of mutation-site side-chain heavy atoms present in both the prediction and experimental mutant; backbone atoms N, C*α*, C, O, and OXT are excluded. After the global fit and the optimal relabeling of chemically symmetry-equivalent atoms, denoted by 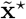, the score is

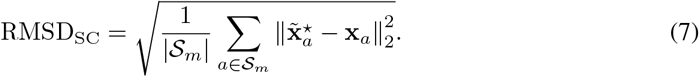

This metric is reported only when at least one comparable side-chain heavy atom is available.

### Mutation-response RMSD_8Å_ (MR-RMSD)

To measure whether a method reproduces the observed reference-to-mutant motion, we independently align the prediction and experimental mutant to the reference over *C*. Let *A*_**w**_(·) denote this Kabsch alignment and define

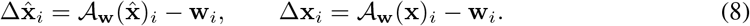

The mutation-response error in the same reference-defined neighborhood is

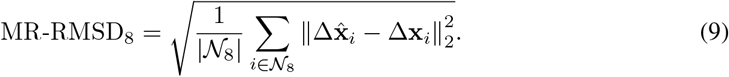

Lower values indicate more accurate prediction of the mutation-induced C*α* displacement field.

For sequence-only predictors, which generate only the mutant structure, MR-RMSD_8Å_ additionally contains ordinary de novo prediction error and is therefore interpreted as a practical reference-relative benchmark score rather than an isolated measure of mutation sensitivity.

